# Spatial profiling of ovarian clear cell carcinoma reveals immune-hot features

**DOI:** 10.1101/2023.07.27.550775

**Authors:** Ya-Ting Tai, Wei-Chou Lin, Duncan Yi-Te Wang, Jieru Ye, Tuan Zea Tan, Lin-Hung Wei, Ruby Yun-Ju Huang

**Affiliations:** Department of Obstetrics & Gynecology, College of Medicine, National Taiwan University, Taipei, Taiwan; Department of Pathology, National Taiwan University Hospital, Taipei, Taiwan; School of Medicine, College of Medicine, National Taiwan University, Taipei, 10051, Taiwan; Cancer Science Institute of Singapore, National University of Singapore, Center for Translational Medicine, Singapore 117599; Graduate Institute of Oncology, College of Medicine, National Taiwan University, Taipei, Taiwan; Department of Obstetrics & Gynecology, Yong Loo Lin School of Medicine, National University of Singapore, Singapore 119077

**Author notes:** Corresponding Author: Professor Ruby Yun-Ju Huang Affiliations: School of Medicine, College of Medicine, National Taiwan University Address: No. 1, Ren-Ai Road Section I, Taipei, 10051, Taiwan.

## Abstract

**Introduction:** OCCC has high incidence in Asia with frequent occurrence at early stage but without sufficient data on molecular stratification for high-risk patients. Recently, immune-hot features have been proposed as an indicator for poor prognosis for early-stage OCCC. Specific patterns of intra-tumoral heterogeneity (ITH) associated with immune-hot features need to be defined.

**Methods:** Formalin-fixed paraffine embedded (FFPE) tumor sections from 10 early-stage OCCC patients were included. Digital Spatial Profiling (DSP) of 18 protein targets was conducted by using the nanoString GeoMx system to profile selected regions of interest (ROIs) based on the reference H&E staining morphology. Areas of illumination (AOIs) were defined according to ROI segmentation by the fluorescence signals of visualization markers pan-cytokeratin (PanCK), CD45, or DNA.

**Results:** Unsupervised hierarchical clustering of 252 AOIs from 229 ROIs showed that PanCK segments expressed different combinations of immune markers suggestive of immune mimicry features. Three immune-hot clusters were identified: granzyme B high (C1-a), immune signal high (C1-b) and immune-like cells (C1-c); two immune-cold clusters were identified: fibronectin-high (C2-a) and signal-cold (C2-b). Immune cells around C1-b and C1-c PanCK+ AOIs were tumor infiltrating immune cells (TIIs) with higher expression of CD68, while those around C1-a, C2-a and C2-b PanCK+ AOIs were non-TIIs with higher expression of SMA. C1-c and C2-a PanCK+ AOIs were associated with OCCC recurrence. TIIs had higher frequencies in C1-b and C1-c PanCK+ AOIs and were associated with OCCC recurrence. Correlating with morphology, tumor samples with recurrence showed higher frequency of papillary pattern. Plus, ROIs with papillary pattern had extremely high frequency of PanCK segments of C1-c feature, higher frequency of TIIs, and macrophage lineage immune mimicry with high intensity of CD68.

**Conclusions:** Spatial profiling of early-stage OCCC tumors revealed that immune mimicry of tumor cells, the presence of TIIs, and papillary pattern in morphology were associated with recurrence.

## Introduction

In 2023, it is estimated that ovarian cancer is the 5^th^ leading cause of cancer mortality in females in the world while not being the top 10 newly diagnosed cancer in population[1]. This features high mortality of ovarian cancer comparing to its relatively low incidence. Epithelial ovarian carcinoma accounts for more than 90% of cases and can be further divided into at least 5 subtypes according to histological and molecular genetics features [2-4]. Among these subtypes, ovarian clear cell carcinoma (OCCC) distinguishes itself from the other subtypes biologically and clinically.

Biologically, OCCC is featured with its complexity of inter-tumoral and intra-tumoral heterogeneity (ITH). Many studies have focused on the molecular and genetic heterogeneity[5, 6] to decipher the mechanisms behind OCCC progression in hope to develop tailored treatments. Previous research in our lab has shown that various gene expression subtypes in OCCC featuring mechanisms such as epithelial-mesenchymal transition (EMT) or immune-related signatures and these would drive tumor progression and correlate with outcomes [7, 8]. These biological features constitute the inter-tumoral heterogeneity of OCCC which would be helpful on patient stratification for potential therapeutic intervention. Clinically, although the majority of OCCC is diagnosed at early stage, metastatic or recurrent OCCC poses great challenges due to limited therapeutic strategies[4, 9]. Therefore, biomarkers are required for helping to identify patients who are at high recurrence risk to design tailored novel treatments. In recent years, the presence of tumor infiltrating lymphocytes (TILs) has been suggested to be a prognostic factor for solid tumors including ovarian carcinoma [10-12]. The prognostic role of TILs in OCCC has not been established and it is not clear whether TILs could serve as a clinical indicator for OCCC recurrence.

OCCC also exists heterogeneity in terms of the pathologic architecture. OCCC has been reported to consist of 3 distinct pathological patterns, papillary, tubulocystic and solid. *Veras et al.* [13] provided some indirect evidence showing the relevance between OCCC’s morphology and the prognosis. Similarly, in clear cell renal cell carcinoma (ccRCC), which shares similar histological morphology with OCCC, *Qi Cai et al.* [14] defined 33 phenotypes according to spatial architecture, cytologic feature and the tumor microenvironment, and illustrated the associations with clinical behaviors such as prognosis and drug resistance. Meanwhile, it is unclear how the complex morphology in OCCC would contribute to the molecular or genetic heterogeneity. Previous studies have focused on the relationship between the OCCC morphology and protein expressions in immunohistochemistry[15, 16]. However, it remains unclear whether these morphological features correlate with other molecular signals and serve as a predictor of disease outcomes. Therefore, it is reasonable to further explore the morphological heterogeneity and the molecular intra-tumoral heterogeneity (ITH) in OCCC.

Digital Spatial Profiling (DSP) is a promising method for the understanding of the spatial distribution of proteins or RNAs within the tissue via advanced multiplexing and quantification of the markers[17-19]. Furthermore, the visualization of the tissue morphology allows users to define region of interest (ROI) and to detect multiple molecular targets *in situ* simultaneously without tissue damage[18, 19]. Hence, in this study, we utilized the DSP technology to gather the spatial information of molecular and morphology heterogeneity and to further identify the geospatial correlation of TILs and other immune cells in early-staged OCCC tumors. We aimed to provide new insights by using the state-of-the-art spatial technology to identify potential biomarkers strategy for OCCC patient stratification.

## Materials and methods

### Clinical cohort and data

An Institutional Review Board (IRB) approved retrospective cohort study (IRB No. 202008022RINB) was performed at the National Taiwan University Hospital (NTUH). A clinical cohort of 195 Stage I/II ovarian clear cell carcinoma (OCCC) patients was reviewed[20]. Ten patients from the cohort were selected for this study to perform Digital Spatial Profiling (DSP). The disease status of these patients is summarized in **Table 1**. The average age of diagnosis was 50.8 years old. All patients underwent adjuvant chemotherapy in NTUH after optimal debulking surgery except the patient of Sample 5, who lost follow-up at the outpatient clinic after optimal debulking surgery (**Supplementary Table 1**). Among the three patients with recurrence, two expired and one lost follow-up and the average progression-free survival (PFS) time for these 3 patients was 4.3 months. The detailed timelines of their disease courses are shown in **Supplementary figure 1**. Upon recurrence, ascites and peritoneal seedings were both found in their first-spotted relapse for these three patients. They received different secondary treatment regimens following recurrence.

**Table 1.** Clinical characteristics of the patients (n=10) Variables Values.

| Variables | Values |
| --- | --- |
| Ages (years) | 50.8 ± 12.6 |
| FIGO stage |  |
| IC1 | 6 (60%) |
| IC2 | 1 (10%) |
| IC3 | 1 (10%) |
| IIA | 1 (10%) |
| IIB | 1 (10%) |
| Recurrence | 3 (30%) |
| Progression free survival (month) | 29.1 ± 22.9 |
| Death | 2 (20%) |

### Sample preparation

Archival formalin-fixed paraffin-embedded (FFPE) samples were retrieved and reviewed by one expert pathologist (W.C. Lin) based on H&E stains to confirm the diagnosis of OCCC. FFPE tissue sections (5 μm) were baked at 60LJ for 1 hour. Deparaffinization was performed using CitriSolv and sections were rehydrated in 100% and 95% ethanol sequentially and followed by wash in ddH2O. Antigen retrieval was performed by boiling at 121LJ in pH 6 Citrate buffer solution for 15 minutes in a pressure cooker. Tissue sections were blocked in blocking buffer for 1 hour at room temperature before incubated with nanoString GeoMx DSP Immune Cell Profiling (a panel of antibodies with UV photocleavable oligonucleotide barcodes), PanCK-Cy3 (1:40) and CD45-Texas Red (1:40) antibodies overnight at 4LJ in dark. Sections were stained in SYTO13 (1:10) for 15 minutes on the following day after post-fixed in 4% paraformaldehyde (PFA).

### Digital Spatial Profiler (DSP)

Formalin-fixed paraffin-embedded (FFPE) samples are incubated with the mixture antibodies of visualization markers (VMs) and DSP probes, which will attach with oligonucleotides by photo-cleavable linkers. Once scanned by DSP, VMs will allow users to select regions of interest (ROIs) and define areas of illumination (AOIs) according to ROI segmentation by pan-cytokeratin (PanCK), CD45 or DNA. UV light projected by DSP machine will then cleave the linkers in those well-defined AOIs, and the released oligonucleotides will be collected by a microcapillary system for further counting by the nCounter system. Selected segmented AOIs from each ROI were profiled by using nanoString Human Protein Core which consisted of 18 protein targets, including Ki-67, fibronectin (FN), PanCK, CTLA4, PD-L1, PD-1, CD20, SMA, Granzyme B (GZMB), CD8, Beta-2-globulin (B2M), CD45, CD11c, HLA-DR, CD4, CD3 and CD56.

### Selection of Region of Interests (ROIs)

A total of 229 ROIs were selected based on their morphology in the H&E stains with 252 areas of illumination (AOIs) collected based on their visualization marker (VM) segmentations. Among them, 209 AOIs were segmented by Pan-cytokeratin (PanCK), 41 AOIs were segmented by CD45, and 2 AOIs were segmented by DNA. 23 ROIs were positive for both PanCK and CD45 segments (**Figure 1a**). In 20 ROIs, the PanCK and CD45 segments were intermingled and were defined to contain tumor-infiltrating immune cells (TIIs). However, in 3 ROIs, PanCK and CD45 segments were geospatially separated, indicating that immune cells just surrounded tumor cells and didn’t infiltrate them (**Figure 1b**). These were defined to contain non-TIIs.

**Fig. 1:**
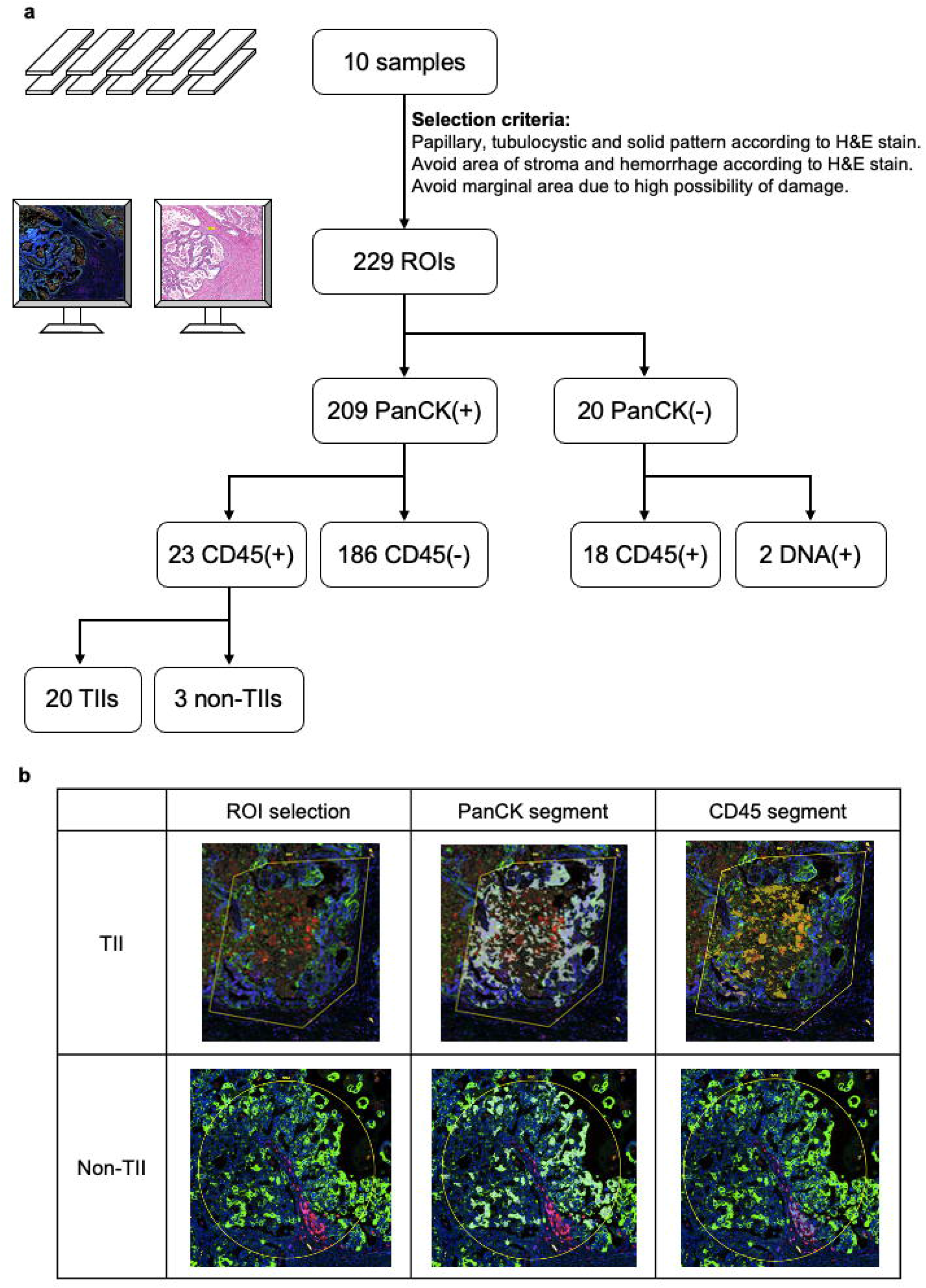
ROI selection. **a,** Workflow of ROI selection. **b,** Identification of TII (upper panels) and non-TII (lower panels). H&E-stained sections were used for morphology confirmation prior to sample preparation. Images of VM-stained fluorescence of DNA (blue), PanCK (green), and CD45 (red) were labeled with selected ROIs.

### Data normalization and visualization

The readouts from nCounter (version 4.0.0.3) were transferred to GeoMx, QC and normalization using the built-in data analysis software (version 2.1.0.33). The raw data was first normalized to ERCC (External RNA Control Consortium) spike-in controls. Subsequently, the spike-in normalized data were scaled by geometric mean of nuclei counts, and by geometric mean of Rb IgG. The normalized data were log-transformed, and genes, arrays were mean-centered by Cluster 3.0[21]. Clustering was also performed using Cluster 3.0 using similarity metric of Pearson correlation, and tree-construction method of centroid linkage. The heatmap of the clustered data was generated with Java Treeview (version 1.1.6r4)[22]. The normalized data were extracted and made into box plots. Statistical significance of association was assessed using Chi-square test whereas mean difference was assessed using ANOVA test.

## Results

### Clustering of AOIs revealed PanCK segments harboring various immune mimicry features

From the unsupervised hierarchical clustering, two major clusters were identified (**Figure 2a**): cluster 1 (C1) as the ‘immune hot’ cluster and cluster 2 (C2) as the ‘immune cold’ cluster. C1 and C2 could be further divided into several sub-clusters. We named PanCK segments from these sub-clusters according to their general shared features. C1-a PanCK+ AOIs were ‘granzyme B (GZMB) high group’, for they represented a group of epithelial cells with high signals of PanCK, CD20, and GZMB. C1-b PanCK+ AOIs were named as ‘immune signal high group’, representing epithelial cells characteristic of high immune signals with equally strong expression of myeloid and lymphoid cell markers. The myeloid signals in this C1-b cluster included CD11c and CD68, and the lymphoid signals included HLA-DR, CD4, CD3, and CD56. C1-c PanCK+ AOIs were ‘immune-like group’ as they showed relatively low expression of PanCK and were co-clustered with CD45+ AOIs. C2-a PanCK+ AOIs were referred as ‘fibronectin (FN) high group’, revealing a group of epithelial cells with high expression levels of FN, PanCK, and myeloid cell marker such as CD68. C2-b PanCK+ AOIs were named as ‘signal cold group’, featuring a group of epithelial cells with relatively low expression of PanCK and relatively high expression of immune checkpoint signals, CTLA4 and PD-1. These spatially resolved PanCK+ segments were found to express different combinations of immune markers suggestive of immune mimicry features.

**Fig. 2:**
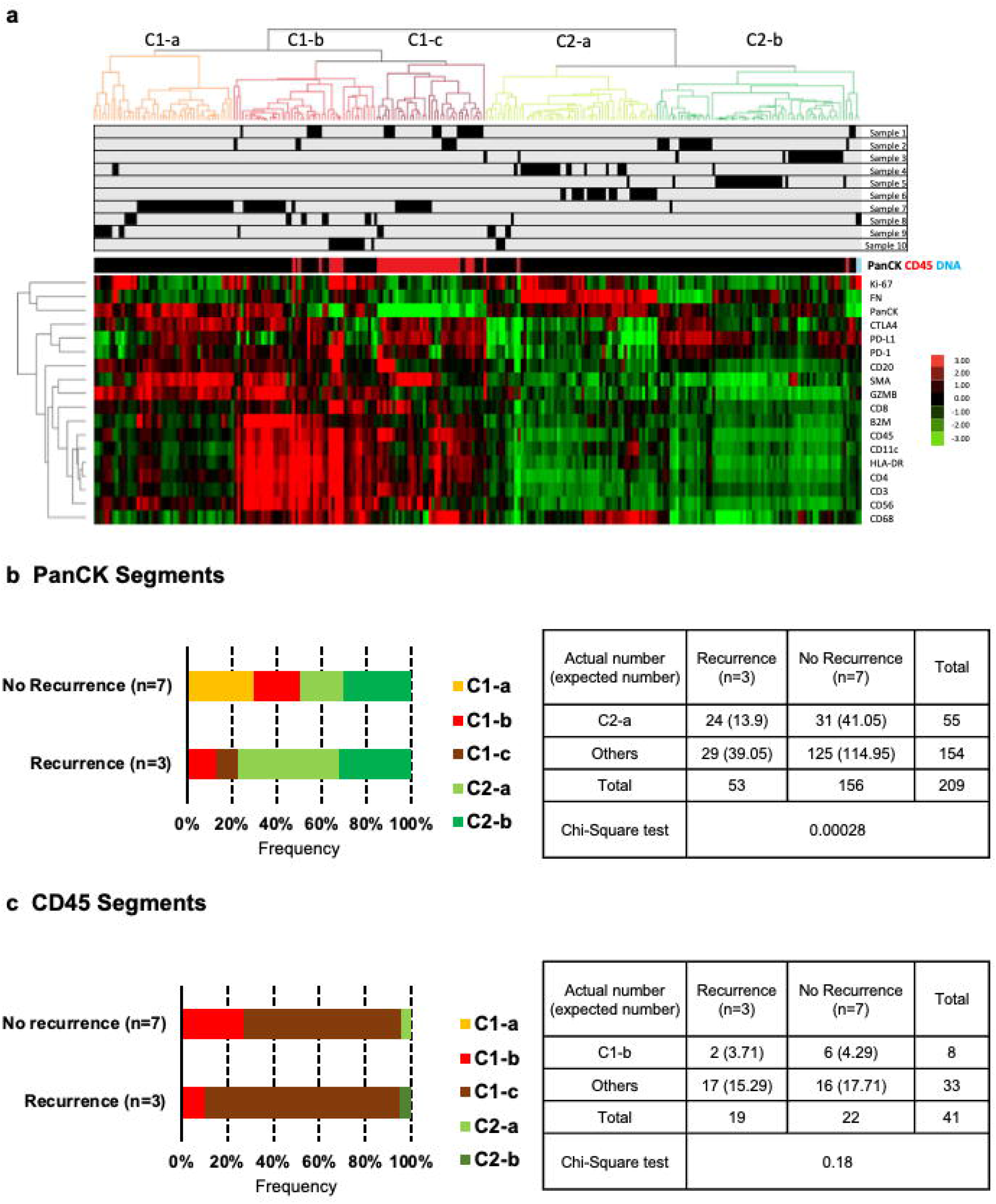
Clustering of AOIs and the association with tumor recurrence. **a,** Unsupervised hierarchical clustering of the 18-plex protein expression data. PanCK-positive segments (black stripes), CD45-positive segments (red stripes) and DNA-positive (blue stripes) among all analyzed AOIs. Subclusters from left to right: C1-a (orange) named as GZMB high group, C1-b (red) named as immune signal high group, C1-c (brown) named as immune-like group, C2-a (light green) named as FN high group, C2-b (dark green) named as signal cold group. Each black stripe in the gray panels indicates one AOI from the specific sample. The heatmap represents the normalized intensity of the 18-plex protein expression. **b,** The proportion of PanCK segments belonging to different subcluster (C1-a, orange; C1-b, red; C1-c, brown; C2-a, light green; C2-b, dark green) in samples with recurrence or with no recurrence. No C1-a (orange) PanCK segment occurred in samples with recurrence. No C1-c (brown) PanCK segment occurred in samples without recurrence. Chi-Square test revealed that C2-a (light green) PanCK segments had higher proportion in samples with recurrence. **c,** The proportion of CD45 segments belonging to different subcluster in samples with recurrence or with no recurrence. Chi-Square test revealed that no specific subcluster of CD45 segments had higher frequency in samples with recurrence.

CD45+ AOIs were found to be grouped within certain clusters. There was no CD45 segment found in C1-a. Majority of the CD45+ AOIs were grouped within the C1-b and C1-c clusters. C1-b CD45+ AOIs were named as ‘signal high immune cells’, for they expressed extremely strong immune-related signals including beta-2-globulin (B2M), CD45, CD11c, HLA-DR, CD4, CD3, and CD56. These signals suggested the presence of antigen presenting cells (APCs), natural killer (NK) cells, and T helper cells. C1-c CD45+ AOIs were named as ‘immune checkpoint signal high immune cells’, for they expressed strong signals of CTLA4, PD-L1 and PD-1. There was only one CD45+ AOI found in C2-a and C2-b, separately.

### The presence of PanCK segments within immune mimicry clusters and tumor-infiltrating immune cells (TIIs) were associated with OCCC recurrence

We noticed that each cluster was contributed by AOIs from different patients indicating the existence of inter-tumoral heterogeneity (**Supplementary Table 1**). We next explored whether these clusters would associate with patients’ outcomes. PanCK segments of C1-a only appeared in patients without recurrence, while those of C1-c only appeared in patients with recurrence (**Figure 2b, Supplementary figure 2**). PanCK segments of C1-b, C2-a, and C2-b clusters were present in patients with and without recurrence but at varying frequencies (**Figure 2b**). C2-a PanCK+ AOIs showed a significantly higher frequency in patients with recurrence (45.3% in recurrence vs 19.9% in no recurrence, Chi-square test: *p* =2.8E-04). These data suggested that tumor cells with C1-c and C2-a features were associated with OCCC recurrence. However, there was no significant difference in the cluster distribution frequency of CD45 segments between tumor samples with recurrence and without recurrence (**Figure 2c, Supplementary figure 2**).

We continued to explore the geospatial relationship between tumor cell clusters and immune cells within the neighborhood microenvironments and its association with patients’ outcomes. Tumor samples from patients with recurrence had higher frequency of TII occurrence (66.7% in recurrence vs 14.3% in no recurrence, Chi-Square test: *p* =0.098) (**Figure 3a, Supplementary figure 2, Supplementary Table 1-3**). TIIs did not appear in PanCK segments of C1-a (0%, Chi-square: *p* =1.56E-05); instead, TIIs appeared in PanCK segments of C1-b and C1-c with higher frequencies (45% in C1-b, Chi-square: *p* =0.046; 25% in C1-c, Chi-square: *p* =0.046) (**Figure 3b, 3c**). We further compared the expression of protein signals from PanCK+ AOIs with TIIs and without TIIs. Compared to TII-PanCK+ AOIs, TII+ PanCK+ AOIs showed higher expression levels of CD45, CTLA4, PD-L1, CD8, CD11c, CD68, HLA-DR, CD3, and CD4 with lower expression levels in PanCK and FN (**Figure 3d, Supplementary figure 3**). This suggested that tumor cells surrounded by TIIs harbored significant immune mimicry features and expressed selected immune markers.

**Fig. 3:**
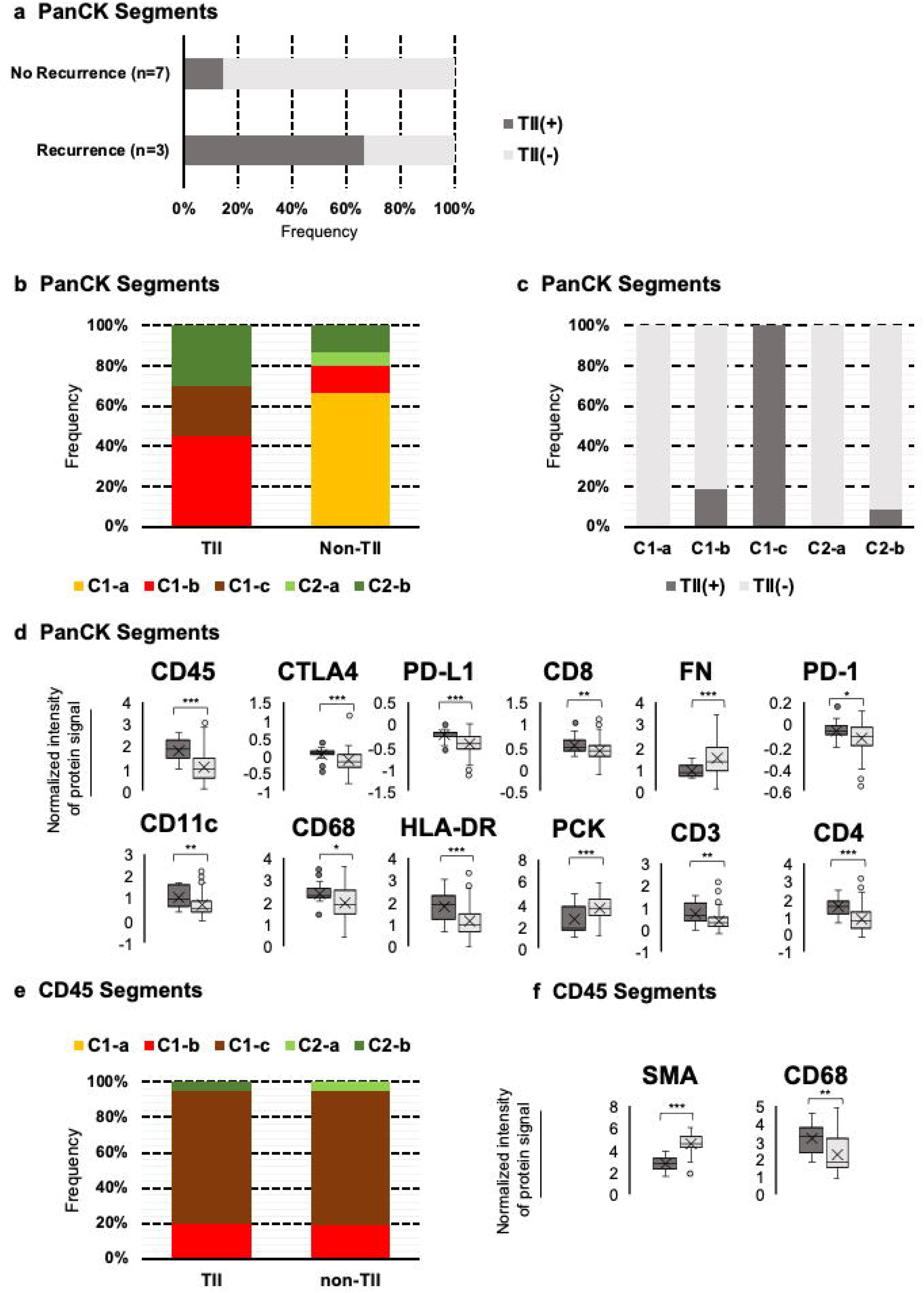
The characteristics of tumor-infiltrating immune cells (TIIs) and the surrounding microenvironment. **a,** The proportion that PanCK segments containing tumor-infiltrating immune cells (TIIs) (light gray) or not (dark gray) in samples with recurrence or with no recurrence. **b,** The proportion that PanCK segments belonging to different subcluster in samples with TIIs and with no TIIs. **c,** The proportion that PanCK segments containing TIIs or not in different subcluster. **d,** The intensity of the normalized protein signals from PanCK segments containing TIIs and those not containing TIIs with statistical significance by one-way ANOVA, **p <0.01, ***p <0.001. Y-axis: numeric values of normalized protein expression level (extracted from the heatmap). **e,** The proportion that different CD45 segments subcluster in the immune cells were classified as TIIs and non-TIIs. **f,** The intensity of the normalized protein signals from CD45 segments of TIIs and non-TIIs with statistical significance by one-way ANOVA, **p <0.01, ***p <0.001. Y-axis: numeric values of normalized protein expression level (extracted from the heatmap).

However, there was no significant difference in cluster distribution of CD45 segments between TIIs and non-TIIs (**Figure 3e**). The expression of protein signals from CD45+ AOIs with TIIs and without TIIs revealed that TIIs showed stronger expression of CD68 (*p* =4.5E-03) and non-TIIs showed stronger expression of SMA (*p* =2.6E-08) (**Figure 3f, Supplementary figure 4**). This indicated that the TIIs could be more enriched of immune cells from the macrophage lineage.

We next explored the immune cells around tumor cells with immune mimicry features. Immune cells around C1-a PanCK+ AOIs were found to be non-TIIs, while those around C1-b and C1-c PanCK+ AOIs were found to be TIIs (**Supplementary figure 2**). Immune cells around C1-a PanCK+ AOIs showed higher protein expression of SMA and lower protein expression of CD68 compared to those around C1-b and C1-c PanCK+ AOIs. Immune cells around C1-b PanCK+ AOIs showed higher protein expression of CD45 comparing to those around C1-a and C1-c PanCK+ AOIs, higher protein expression of CD3, CD4 and CD11c comparing to those around C1-a PanCK+ AOIs and higher protein expression of GZMB comparing to those around C1-c PanCK+ AOIs (**Supplementary figure 5**). This suggested that tumor cells with ‘GZMB high’ feature would more likely attract non-TIIs with high expression of SMA. Tumor cells with ‘immune signal high’ feature would more likely attract myeloid and lymphoid lineage of infiltrating immune cells with high expression of CD45, CD3, CD4, GZMB, CD11c. Tumor cells with ‘immune-like’ feature would more likely attract macrophages with high expression of CD68 while relatively low expression of CD3, CD20 and CD8.

### Cluster mixture of PanCK segments and the presence of TIIs

There was significant intertumoral and intratumoral heterogeneity of the number of different clusters in cluster mixture (**Supplementary figure 2, 6a**). Based on the cluster annotations of PanCK segments, 2 (20%) tumors showed high homogeneity of PanCK segments with the presence of only one cluster. These two homogeneous tumors, sample 3 and 6, were composed of the immune cold group, C2-b and C2-a, respectively. No TII was found in the two homogeneous tumors with immune cold PanCK clusters (**Supplementary figure 2, 6a**).

The other 8 (80%) tumors showed heterogeneity in the cluster mixture of PanCK segments with 3 (30%) tumors mixed by two clusters and 5 (50%) mixed by three clusters. The 2-cluster and 3-cluster heterogeneous tumors showed the composition of immune cold and immune hot clusters without significant pattern of mixture. In addition, there was no significant difference between tumors from patients with or without recurrence in terms of the number of cluster mixtures (**Supplementary figure 6b**). In these heterogeneous tumors, TIIs were only found in 37.5% (3/8, sample 2, sample 10, sample 1) of tumors (**Supplementary figure 2, 6a**). In 2-cluster tumors with TIIs (2/3, 66.7%), the TIIs were adjacent to the C1-b PanCK+ AOIs. In 3-cluster tumors, TIIs were present in sample 1 which was the only one present with TIIs (1/5, 20%) and was the only one which did not comprise of the C1-a cluster. TIIs were not present in either of the other four samples containing C1-a cluster in the cluster mixture (**Supplementary figure 2, 6a**). In sum, the number of cluster mixture did not correlate with the presence of TIIs nor recurrence in OCCC. It was the nature of cluster mixture (eg. the presence of C1-a cluster) which might determine the presence of TIIs.

### Intratumoral heterogeneity of morphology is associated with OCCC recurrence and TII frequency

There are various degrees of heterogeneity of morphology among different tumor samples and within each tumor sample (**Supplementary figure 2, 7a**). We noticed that tumor samples with recurrence, compared to those without recurrence, showed higher frequency (54.7%, Chi-Square test: p=1.01E-04) of the papillary pattern and lower frequency (18.9%, Chi-Square test: p=7.21E-03) of the tubulocystic pattern (**Figure 4a**). We then investigated the correlation between tumor morphology and TME of the clusters (**Supplementary Table 4**). ROIs with a papillary pattern showed extremely low frequency (6.2%, Chi-Square test: p=2.01E-04) of C1-a PanCK segments, and ROIs with a solid pattern showed extremely low frequency (2.8%, Chi-Square test: p=1.93E-05) of C1-b. Interestingly, C1-c PanCK segments only appeared in those ROIs with a papillary pattern (**Figure 4b, 4c**). These data corroborated with the significant exclusiveness found between recurrence and clusters (**Figure 2b**) and suggested that morphological heterogeneity might be a prognostic indicator in OCCC. The expression of the 18-plex protein signals among different morphologies was thus further explored (**Supplementary figure 7b**). The papillary pattern showed low expression of PanCK and high expression of CD68 with statistical significance. The tubulocystic pattern showed high expression of SMA, CD20, FN and GZMB with statistical significance. The solid pattern showed low expression of immune signals with statistical significance including CD45, CD11c, HLA-DR, CD3, CD4 and CD56, confirming the relatively cold immune nature.

**Fig. 4:**
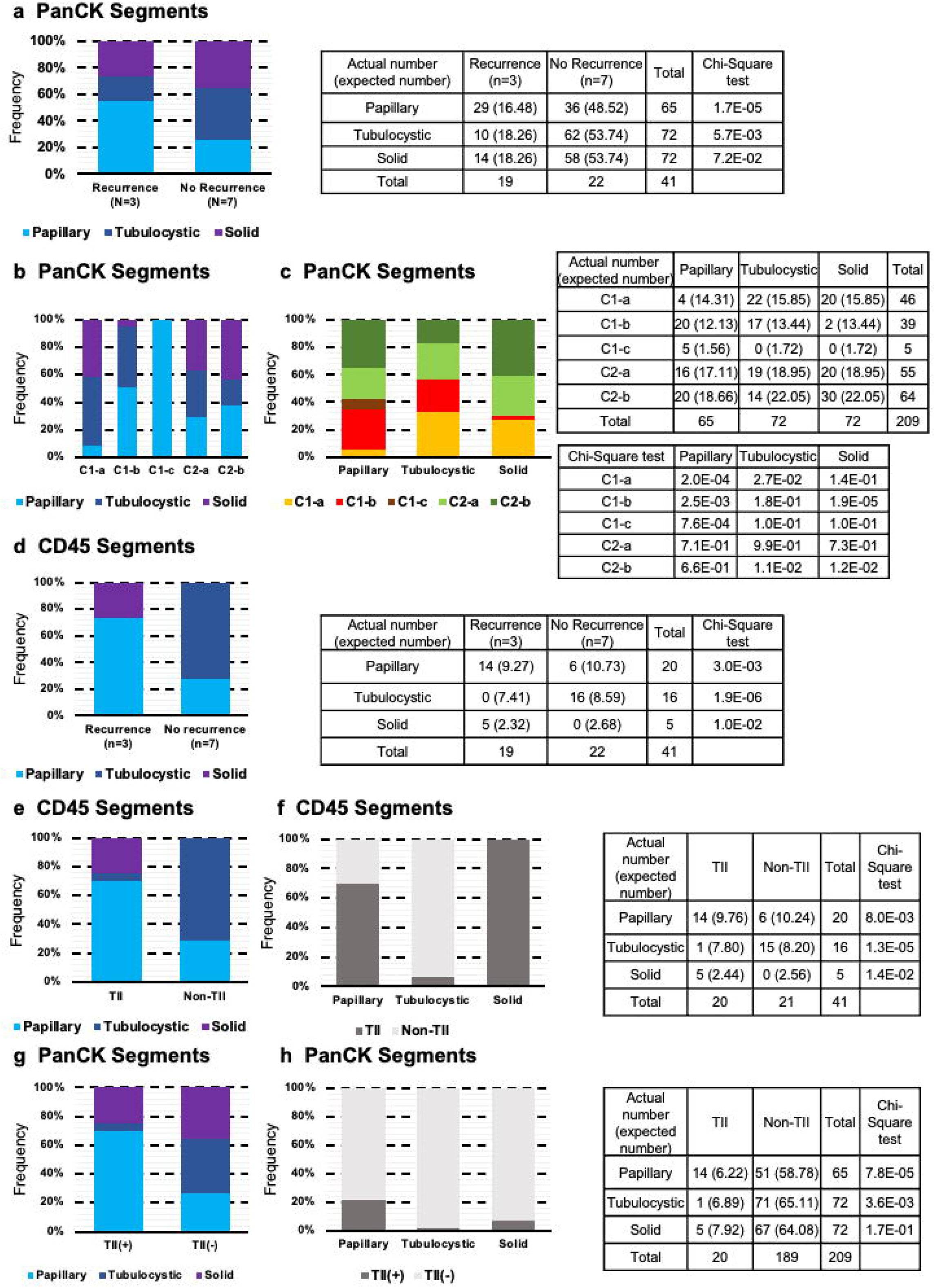
Tumor morphology and the association with tumor recurrence and TIIs. **a,** The proportion that PanCK segments were identified as papillary patterns (light blue), tubulocystic patterns (dark blue), and solid patterns (purple) in samples with recurrence or with no recurrence. **b,** The proportion that PanCK segments identified as different morphological patterns in different subcluster. **c,** The proportion that PanCK segments belonging to different subcluster in different morphological patterns. **d,** The proportion that CD45 segments were identified as different morphological patterns in samples with recurrence or with no recurrence. **e,** The proportion that CD45 segments identified as different morphological patterns in samples with TIIs and with no TIIs. **f,** The proportion that CD45 segments containing TIIs or not in different morphological patterns. **g,** The proportion that PanCK segments identified as different morphological patterns in samples with TIIs and with no TIIs. **h,** The proportion that PanCK segments containing TIIs or not in different morphological patterns.

The intertumoral and intratumoral heterogeneity of morphology in CD45+ AOIs were also noted (**Supplementary figure 2, Supplementary Table 5**). Intriguingly, immune cells within the papillary ROIs had higher frequency (73.7%, Chi-Square test: p=3.03E-03) in tumor samples with recurrence (**Figure 4d**). In contrast, those within the tubulocystic ROIs only occurred in tumor samples without recurrence (**Figure 4d**). Immune cells occurred with lower frequency (12.2%) within the solid pattern, and these immune cells within the solid pattern only occurred in tumor samples with recurrence. This prompted us to examine the correlation between the presence of TIIs and the morphology patterns (**Supplementary Table 5**). Compared to non-TII CD45+ AOIs, TIIs showed higher frequency in the papillary pattern (70.0%, Chi-Square test: p=7.98E-03) and lower frequency in the tubulocystic pattern (5.0%, Chi-Square test: p=1.31E-05) (**Figure 4e, 4f**). The immune cells present in the tubulocystic pattern were almost exclusively non-TIIs (**Figure 4e, 4f**). TIIs were found with low frequency (5.0%, Chi-Square test: p=3.56E-03) in PanCK segments with a tubulocystic pattern and with high frequency (70.0%, Chi-Square test: p=7.75E-05) in PanCK segments with papillary pattern (**Figure 4g, 4h**). The data clearly demonstrated significant correlation between OCCC recurrence with the presence of TIIs within the papillary morphology of the PanCK segments harboring immune hot cluster features.

## Discussion

Immune-related subtypes have been identified to correlate with patient outcomes in OCCC [7, 20, 23]. These studies were based on bulk analysis of tumor samples and thus provided the perspective of intertumoral heterogeneity in OCCC. In this study, by using digital spatial profiling (DSP) technology, we explore the intra-tumoral heterogeneity (ITH) of 10 OCCC tumors and analyzed the expression patterns of molecular signals in relation to the geospatial context and the pathological morphology. We identified 5 clusters of OCCC cells with immune mimicry using spatially resolved technique, which could provide clear compartmentalization between tumor cells and immune cells. Immune mimicry in tumor cells has been proposed to be a mechanism of immune evasion [24-26], and Gao et al. [27] has demonstrated that various immune mimicry of tumor cells could lead to different clinical outcomes. There are 3 types of immune hot features indicative of immune mimicry of tumor cells. OCCC with ‘GZMB high’ epithelial cells had favorable prognosis, in which tumor cells featuring high level of granzyme B and CD20 attracted non-TII featuring high level of SMA, which echoed with previous studies suggesting that tumor-infiltrating CD20+ B cells correlated with favorable prognosis in non-small cell lung, breast, cervical and ovarian cancer[28-31]. Interestingly, this group of epithelial cells expressing high levels of granzyme B and CD56 showed low levels of CD4 and CD8, which indicated the mimicry towards NK cell instead of T cell. Furthermore, non-TIIs featuring high level of SMA were found with higher frequency around this group of epithelial cells. In sum, we supposed that NK cell/B cell mimicry of tumor cells with the co-existing immune cells contributes towards a better TME and lead to favorable prognosis. On the other side, OCCC with ‘immune signal high’ epithelial cells may have poorer prognosis, in which tumor cells were prone to attract TII featuring high level of CD45. The immune signals featuring in this group of tumor cells showed non-specific immune mimicry towards lineages. OCCC with ‘immune-like’ epithelial cells had unfavorable prognosis, in which tumor cells featuring high level of CTLA4 attracting TIIs featuring high level of CD68. We figured that immune checkpoint mimicry of tumor cells could attract antigen presenting cells to elicit immune responses. In this case, the inhibitory immune signals from CTLA4 immune mimicry further induce the macrophage infiltration to contribute to the least permissive TME. Our findings suggest that immune mimicry is a double-edged sword depending on the type of immune masquerade the tumor cells put on and these immune mimicry features provide additional information on the intricacy of the immune landscape within OCCC.

We also noted that tumor samples with ‘FN high’ epithelial cells had poorer outcome indicating that phenotype hybrid OCCC with high level of both PanCK and FN would lead to poorer prognosis. However, these ‘FN high’ tumor cells were relatively cold in immune mimicry expressing scanty myeloid lineage markers and did not attract TIIs. Since FN is a well-documented mesenchymal marker, it is interesting that TII didn’t exist within such mesenchymal-like tumor cells. This agrees with the known immunosuppressive function elicited by EMT [8, 32, 33].

The immune subtype identified by Ye et al., [23] and the Immune-Hot subtype identified by Huang et al., [20] both revealed consistently the adverse tumor biology related to immune features. However, the bulk analysis of these two studies could not delineate whether these immune subtypes were contributed by the infiltration of immune cells or the immune mimicry of tumor cells. The spatially resolved analysis from this current study has shown that infiltration of immune cells and immune mimicry of tumor cells both play crucial roles in shaping the immune subtype in OCCC.

We discovered different combinations of the 5 clusters among OCCC tumors. Intriguingly, epithelial cells with immune hot features did not exist as standalone single cluster. The immune mimicry tumor cells must exist in tumors with certain degree of intra-tumoral heterogeneity, which may imply that these immune mimicry tumor cells were acquired from selection pressure. Our study echoed previous research of tumor immunoscore [34-36] suggesting that the immune process is not always associated with favorable clinical outcomes and could be either tumor-suppressing or tumor-promoting. The cluster mixture containing ‘GZMB high’ epithelial cells could be tumor-suppressing, and that containing ‘immune-like’ epithelial cells could be tumor-promoting.

By re-analyzing normalized data from DSP, we found that tumor cells within the papillary pattern and those with TIIs both featured with low expression of PanCK and high expression of CD68. The TIIs within these tumor cells featured relatively high expression of CD68 themselves. In contrast, higher levels of SMA were found in tumor cells in tubulocystic pattern and immune cells of non-TIIs. Tumors with tubulocystic pattern may be more mesenchymal and disallowed the infiltration of immune cells, and thus had better prognosis. Veras et al. [7] provided some indirect evidence showing the relevance between OCCC’s morphology and the prognosis. They suggested that OCCC could be divided into two categories --- arising from a cyst or from an adenofibroma, which harbored different clinico-pathologic factors including stage at presentation, association with endometriosis, histologic patterns, and survival. Cystic OCCC with the dominant papillary morphology had better prognosis than adenofibromatous OCCC with the dominant tubulocystic morphology. However, there was still more than 50% of OCCC in this study with no predominance of any architectural patterns. Instead, those cases displayed a combination of the 3 patterns. Similarly, in clear cell renal cell carcinoma (ccRCC), which shares similar histological morphology with OCCC, Qi Cai et al. [8] defined 33 phenotypes according to spatial architecture, cytologic feature and the tumor microenvironment, and illustrated the associations with clinical behaviors such as prognosis and drug resistance. After controlling for nucleolar grade and aggressive cytologic feature of sarcomatoid/ rhabdoid elements, 4 aggressive morphologic patterns including tubular component, ChRCC-like pattern, infiltration into the renal parenchyma and necrosis were identified to be predictive of disease-free survival. Both studies suggested that morphology in OCCC may have correlation with clinical outcome, while our study further demonstrated that morphology in OCCC had correlation with the presence of TIIs and clinical outcome.

OCCC is known for its relative resistance to platinum-based chemotherapy, and this might explain its relatively poor prognosis of ovarian carcinoma[37-39]. Therefore, novel treatment options are needed for OCCC. Immunotherapy has been considered to have potential benefits on OCCC patients. However, one should be mindful about patient selection due to the limited efficacy observed in a small number of cases[40, 41]. Biomarkers predictive of therapeutic efficacy have not been identified from a case series of OCCC patients receiving immune checkpoint blockade [42]. However, these patients were in recurrent or advanced diseases and data on the responses to immune checkpoint blockade in the front-line is currently not available. Our study focused on TME of primary OCCC patients, and the findings could suggest potential biomarkers for patient selection of immunotherapy including immune-hot features, TIIs and tumor morphology. The interaction between immune mimicry of tumor cells and the TIIs could be a complex issue, and further investigation is highly warranted.

## Acknowledgements

This work was supported by the Yushan Scholar Program by the Ministry of Education, Taiwan (NTU-112V1402-5) and NTU Core Consortiums (NTUCC-112L894903) to R.Y.-J.H; NTUCC-112L894901 to Lin-Hung Wei.

## Author contributions statement

Conceptualization, R.Y.-J.H. and Lin-Hung Wei; writing, Duncan Yi-Te Wang, Ya-Ting Tai, Wei-Chou Lin and R.Y.-J.H.; DSP data acquisition, Ya-Ting Tai, Jieru Ye and Duncan Yi-Te Wang; data analysis and interpretation, Ya-Ting Tai, Duncan Yi-Te Wang and Tuan Zea Tan; pathology review, Wei-Chou Lin; clinical review, Lin-Hung Wei and Ya-Ting Tai. All authors have read and agreed to the published version of the manuscript.

## Supplementary Figure Legends

**Supplementary figure 1. Treatments on the patients with recurrence**

**a-c,** Timeline of treatment course received by the patient of sample 1, 2 and 6 respectively.

**Supplementary figure 2. AOI distribution of clustering in each sample**

Patients with recurrence: Sample 1,2 and 6. Patients with no recurrence: exclusively. Each sample from left to right: image of visualized markers by DSP and selected ROIs, outline of the morphology and selected AOIs segmented by PanCK, outline of the morphology and selected AOIs segmented by CD45 (while by *DNA in sample 8), image of H&E stain and selected ROIs. The color of each AOI corresponds with the color of cluster in the heatmap. Scale bars indicate 4000 µm.

**Supplementary figure 3. Protein signature of OCCC with TIIs**

The intensity of the normalized protein signals from PanCK segments containing TIIs and those not containing TIIs. Y-axis: numeric values of normalized protein expression level (extracted from the heatmap). One-way ANOVA, *p<0.05, **p <0.01, ***p <0.001.

**Supplementary figure 4. Protein signature of TIIs**

The intensity of the normalized protein signals from CD45 segments of TIIs and non-TIIs. Y-axis: numeric values of normalized protein expression level (extracted from the heatmap). One-way ANOVA, *p<0.05, **p <0.01, ***p <0.001.

**Supplementary figure 5. Immune cells around tumor cells with immune mimicry features**

The intensity of the normalized protein signals from CD45 segments around C1-a (orange), C1-b (red) and C1-c (brown) PanCK+ AOIs. Y-axis: numeric values of normalized protein expression level (extracted from the heatmap). One-way ANOVA, *p<0.05, **p <0.01, ***p <0.001.

**Supplementary figure 6. Intertumoral and intratumoral heterogeneity of cluster mixture**

**a,** The proportion of PanCK segments belonging to different subcluster (C1-a, orange; C1-b, red; C1-c, brown; C2-a, light green; C2-b, dark green) in each sample. Tumor-infiltrating immune cells (TIIs) exist in Sample 1, 2 and 10. **b,** The proportion that samples of different PanCK cluster mixtures (1-cluster, black bar with dots; 2-cluster samples, gray bar with dots; 3-cluster samples, white bar with dots) with recurrence or with no recurrence.

**Supplementary figure 7. Intertumoral and intratumoral heterogeneity of morphology**

**a,** The proportion of PanCK segments with different morphology (papillary pattern, light blue; tubulocystic pattern, dark blue; solid pattern, purple) in each sample. Tumor-infiltrating immune cells (TIIs) exist in Sample 1, 2 and 10. **b,** The intensity of the normalized protein signals from PanCK segments with different morphology. Y-axis: numeric values of normalized protein expression level (extracted from the heatmap). One-way ANOVA, **p <0.01, ***p <0.001.

