## Supplementary Table 1 for "Spatial profiling of ovarian clear cell carcinoma reveals immune-hot features"

Supplementary Table 1. Disease status (till the closure of this study) of each patient of the sample.

NED: no evidence of disease; NA: not available

| Sample | Diagnosis age | Stage | Disease status | Progression free survival |
| --- | --- | --- | --- | --- |
| Sample 1 | 38 | stage IC1,  pT1cN0Mx, right | Death,  survived for 2y9m | 11m |
| Sample 2 | 56 | stage IC2,  pT1c2N0M0, left | Loss of follow up | 17m |
| Sample 3 | 64 | stage IC1,  pT1c1N0Mx, right | NED | No event |
| Sample 4 | 60 | stage IC1,  pT1c1N0Mx, right | NED | No event |
| Sample 5 | 52 | stage IC1,  pT1cN0Mx, right | Loss of F/U | NA |
| Sample 6 | 57 | stage IC3,  pT1c2N0Mx, right | Death,  survived for 2y5m | 12m |
| Sample 7 | 59 | stage IC1,  pT1c1N0M0, left | NED | No event |
| Sample 8 | 46 | Stage IC1,  pT1c1N0M0, right | NED | No event |
| Sample 9 | 54 | stage IIA,  pT2a(m)N0M0, right | NED | No event |
| Sample 10 | 22 | stage IIB,  cT2bN0M0, left | NED | No event |
