## Supplementary Table 2 for "Spatial profiling of ovarian clear cell carcinoma reveals immune-hot features"

Supplementary Table 2. AOI distribution of clustering in each sample.

| Sample 10 | 0 | 0 | 0 (0%) | 8 | 5 | 13 (81.3%) | 0 | 0 | 0 (0%) | 3 | 0 | 3 (18.7%) | 0 | 0 | 0 | 0 (0%) |
| --- | --- | --- | --- | --- | --- | --- | --- | --- | --- | --- | --- | --- | --- | --- | --- | --- |
| Sample 9 | 8 | 0 | 8 (50.0%) | 1 | 0 | 1 (6.3%) | 0 | 2 | 2 (12.5%) | 5 | 0 | 5 (31.2%) | 0 | 0 | 0 | 0 (0%) |
| Sample 8 | 4 | 0 | 4 (25%) | 9 | 0 | 9 (56.3%) | 0 | 0 | 0 (0%) | 1 | 0 | 1 (6.3%) | 0 | 0 | 2 | 2 (12.5%) |
| Sample 7 | 32 | 0 | 32 (53.3%) | 14 | 1 | 15 (25.0%) | 0 | 12 | 12 (20.0%) | 0 | 0 | 0 (0%) | 1 | 0 | 0 | 1 (1.7%) |
| Sample 6 | 0 | 0 | 0 (0%) | 0 | 0 | 0 (0%) | 0 | 0 | 0 (0%) | 24 | 0 | 24 (100%) | 0 | 0 | 0 | 0 (0%) |
| Sample 5 | 0 | 0 | 0 (0%) | 0 | 0 | 0 (0%) | 0 | 0 | 0 (0%) | 1 | 0 | 1 (3.8%) | 25 | 0 | 0 | 25 (96.2%) |
| Sample 4 | 2 | 0 | 2 (8.3%) | 0 | 0 | 0 (0%) | 0 | 0 | 0 (0%) | 21 | 0 | 21 (87.5%) | 1 | 0 | 0 | 1 (4.2%) |
| Sample 3 | 0 | 0 | 0 (0%) | 0 | 0 | 0 (0%) | 0 | 1 | 1 (4.5%) | 0 | 1 | 1 (4.5%) | 20 | 0 | 0 | 20 (90.9%) |
| Sample 2 | 0 | 0 | 0 (0%) | 2 | 1 | 3 (12.5%) | 0 | 5 | 5 (20.8%) | 0 | 0 | 0 (0%) | 15 | 1 | 0 | 16 (66.7%) |
| Sample 1 | 0 | 0 | 0 (0%) | 5 | 1 | 6 (25.0%) | 5 | 11 | 16 (66.7%) | 0 | 0 | 0 (0%) | 2 | 0 | 0 | 2 (8.3%) |
| AOIs | PanCK | CD45 | Total | PanCK | CD45 | Total | PanCK | CD45 | Total | PanCK | CD45 | Total | PanCK | CD45 | DNA | Total |
|  | C1-a | | | C1-b | | | C1-c | | | C2-a | | | C2-b | | | |
