## Supplementary Table 3 for "Spatial profiling of ovarian clear cell carcinoma reveals immune-hot features"

Supplementary Table 3. Distribution of ROI morphology of each sample. The amounts of ROIs with tumor-infiltrating immune cells (TIIs) were as followed: eleven of Sample 1, seven of Sample 2, and two of Sample 10. The two of Sample 3 and the one of Sample 10 were not TIIs.

|  | Papillary | | | Tubulocystic | | | Solid | | | |
| --- | --- | --- | --- | --- | --- | --- | --- | --- | --- | --- |
|  | ROI | AOI | | ROI | AOI | | ROI | AOI | | |
|  |  | PanCK | CD45 |  | PanCK | CD45 |  | PanCK | CD45 | DNA |
| Sample 1 | 13*  (100.0%) | 12* | 12* | 0  (0%) | 0 | 0 | 0  (0%) | 0 | 0 | 0 |
| Sample 2 | 6  (35.3%) | 6 | 1 | 2 (11.8%) | 2 | 0 | 9 (52.9%) | 9 | 6 | 0 |
| Sample 3 | 10  (50.0%) | 10 | 1 | 6 (30.0%) | 6 | 1 | 4 (20.0%) | 4 | 0 | 0 |
| Sample 4 | 0  (0%) | 0 | 0 | 14 (58.3%) | 14 | 0 | 10 (41.6%) | 10 | 0 | 0 |
| Sample 5 | 7  (26.9%) | 7 | 0 | 3 (11.5%) | 3 | 0 | 16 (61.5%) | 16 | 0 | 0 |
| Sample 6 | 11  (45.9%) | 11 | 0 | 8 (33.3%) | 8 | 0 | 5 (20.8%) | 5 | 0 | 0 |
| Sample 7 | 0  (0%) | 0 | 0 | 46 (76.7%) | 33 | 13 | 14 (23.3%) | 14 | 0 | 0 |
| Sample 8 | 12  (75.0%) | 12 | 0 | 0 (0%) | 0 | 0 | 4 (25.0%) | 2 | 0 | 2 |
| Sample 9 | 3  (18.8%) | 2 | 1 | 4 (25.0%) | 3 | 1 | 9 (56.2%) | 9 | 0 | 0 |
| Sample 10 | 11  (84.6%) | 9 | 4 | 2 (15.4%) | 2 | 1 | 0  (0%) | 0 | 0 | 0 |
