## Supplementary Table 4 for "Spatial profiling of ovarian clear cell carcinoma reveals immune-hot features"

Supplementary Table 4. Distribution of pathological patterns of ROIs segmented by PanCK in each cluster.

|  | Papillary (n, %) | Tubulocystic (n, %) | Solid (n, %) |
| --- | --- | --- | --- |
| Total | 65, 100.0% | 72, 100.0% | 72, 100.0% |
| C1-a | 4, 6.2% | 22, 30.6% | 20, 27.8% |
| C1-b | 20, 30.8% | 17, 23.6% | 2, 2.8% |
| C1-c | 5, 7.7% | 0, 0.0% | 0, 0.0% |
| C2-a | 16, 24.6% | 19, 26.4% | 20, 27.8% |
| C2-b | 20, 30.8% | 14, 19.4% | 30, 41.7% |
