## Supplementary Table 5 for "Spatial profiling of ovarian clear cell carcinoma reveals immune-hot features"

Supplementary Table 5. Distribution pathological patterns of ROIs segmented by CD45 in each cluster.

| CD45 | Papillary (n, %) | | Tubulocystic (n, %) | | Solid (n, %) | |
| --- | --- | --- | --- | --- | --- | --- |
|  | TIIs | Non-TIIs | TIIs | Non-TIIs | TIIs | Non-TIIs |
| C1-a | 0 | 0 | 0 | 0 | 0 | 0 |
| C1-b | 3 | 3 | 1 | 1 | 0 | 0 |
| C1-c | 10 | 2 | 0 | 14 | 5 | 0 |
| C2-a | 0 | 1 | 0 | 0 | 0 | 0 |
| C2-b | 1 | 0 | 0 | 0 | 0 | 0 |
