## Supplementary figure 1 for "Spatial profiling of ovarian clear cell carcinoma reveals immune-hot features"

a

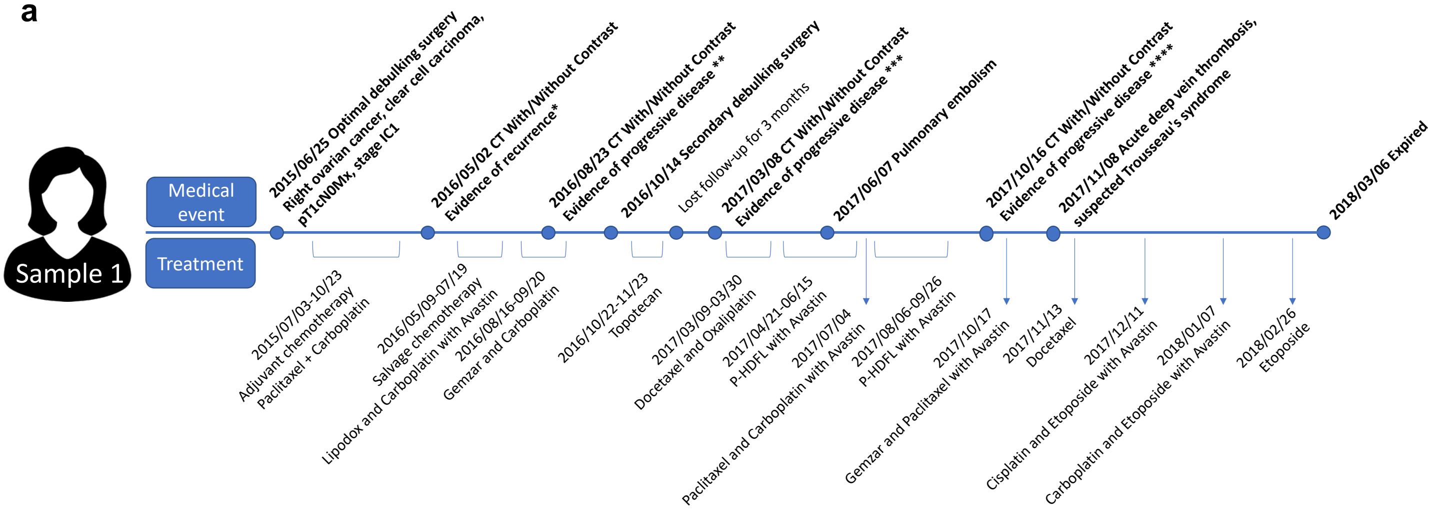


* Ascites and peritoneal carcinomatosis.

** Smaller multiple peritoneal seeding tumors at the perihepatic region and bilateral abdomen

*** Larger peritoneal metastasis and lymphadenopathy at the para-aortic/para-caval region and hepatic hilum

**** 1. Pulmonary emboli in the left tributary, partially regressed
2. Enlarged bilateral mediastinal and axillary lymphadenopathy
3. Multiple peritoneal metastasis with large ascites
4. Peritoneal metastasis invades into the liver through capsule;
5. Lymphadenopathy at the hepatic hilum, para-aortic/para-caval region, stationary
6. Thrombus in the right common/superficial femoral vein

b

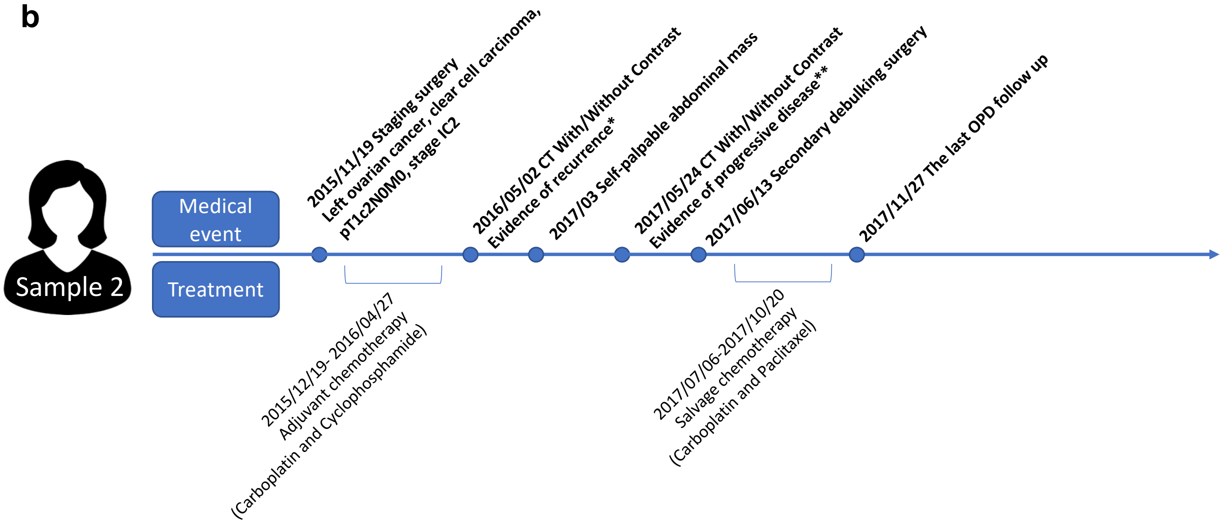


* Much ascites and peritoneal carcinomatosis

**1. Recurrent tumor at the left iliacus muscle and left abdomen about 16cm, new.
2. Lymphadenopathy at the paraaortic region, new.

c

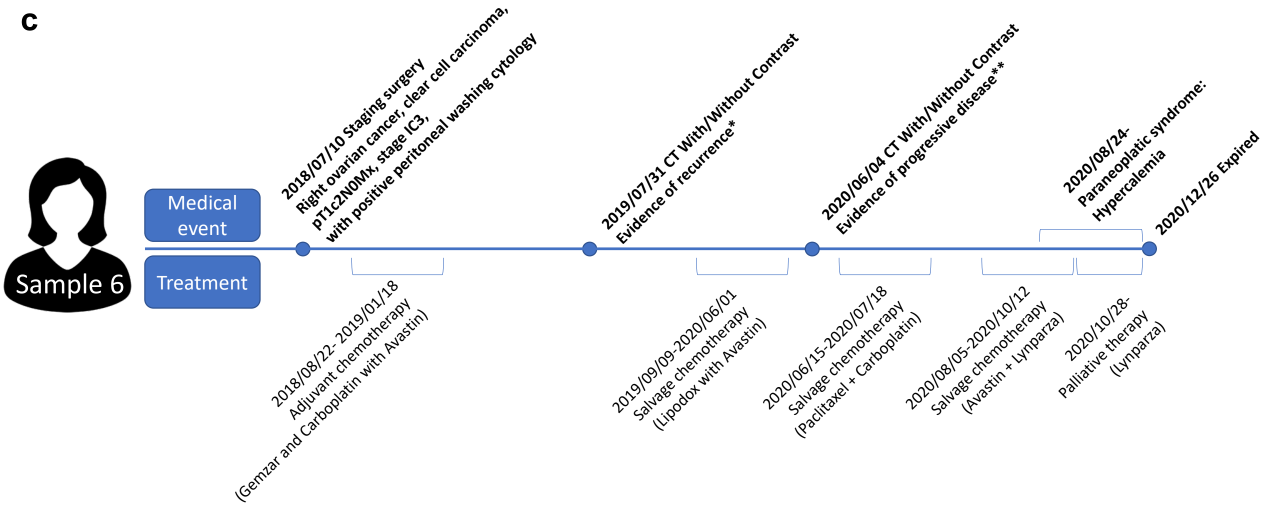


* 1.Right ovarian cancer, status post staging surgery.
2.Small nodules at right subphrenic region, with mild ascites in the pelvic cavity, peritoneal tumor metastasis is suspected.
3.Right pleural effusion with partial atelectasis of right lower lung. Mild pleural thickening, malignant pleural effusion is suspected.
4.One small hemangioma at inferior tip of right hepatic lobe, stable.
**Peritoneal and pleural metastases .
