## Supplementary figures and images for "Spatial profiling of ovarian clear cell carcinoma reveals immune-hot features"

### Supplementary figure 2

#### Supplementary Figure 2

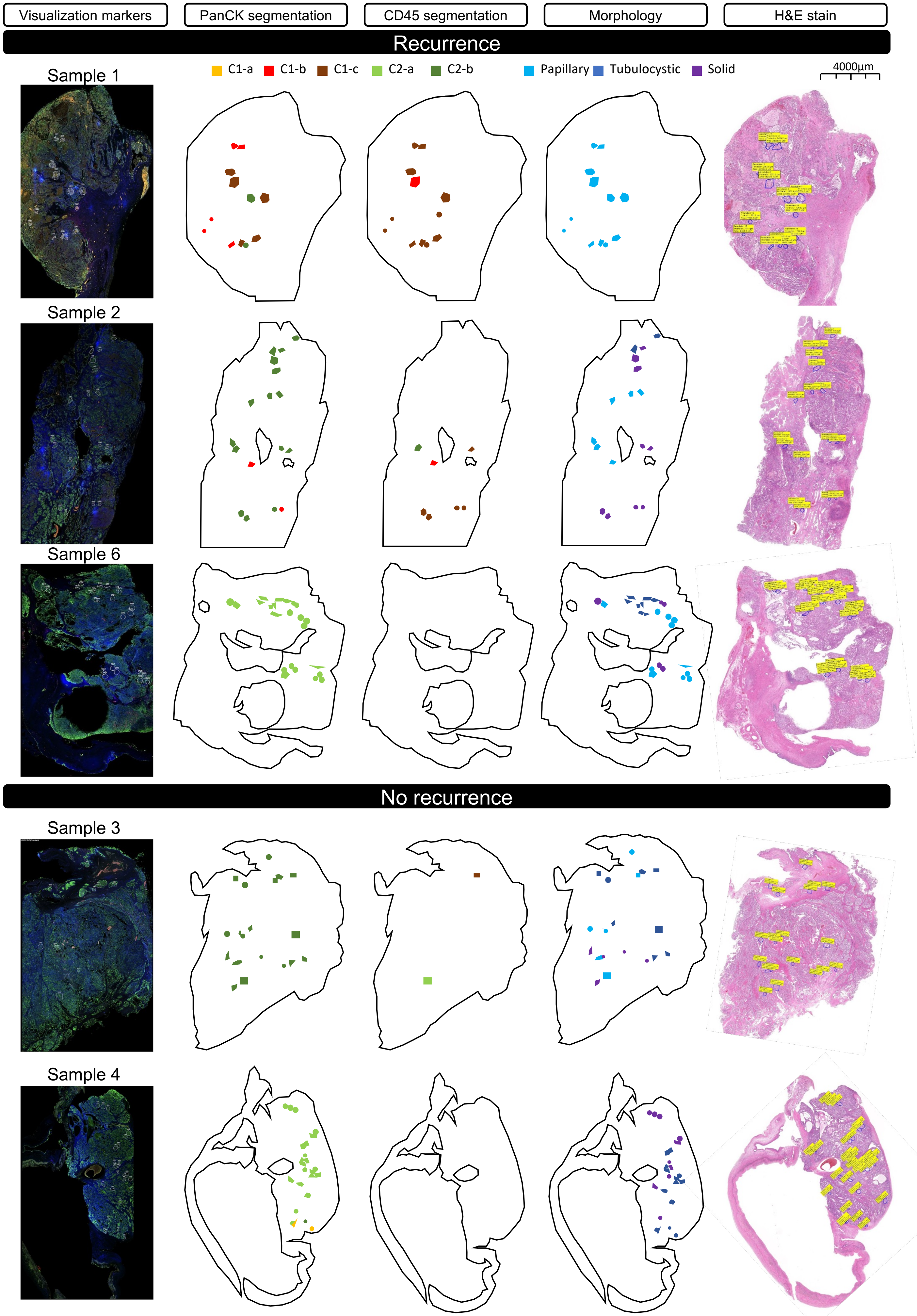

#### Supplementary Figure 2 (continued)

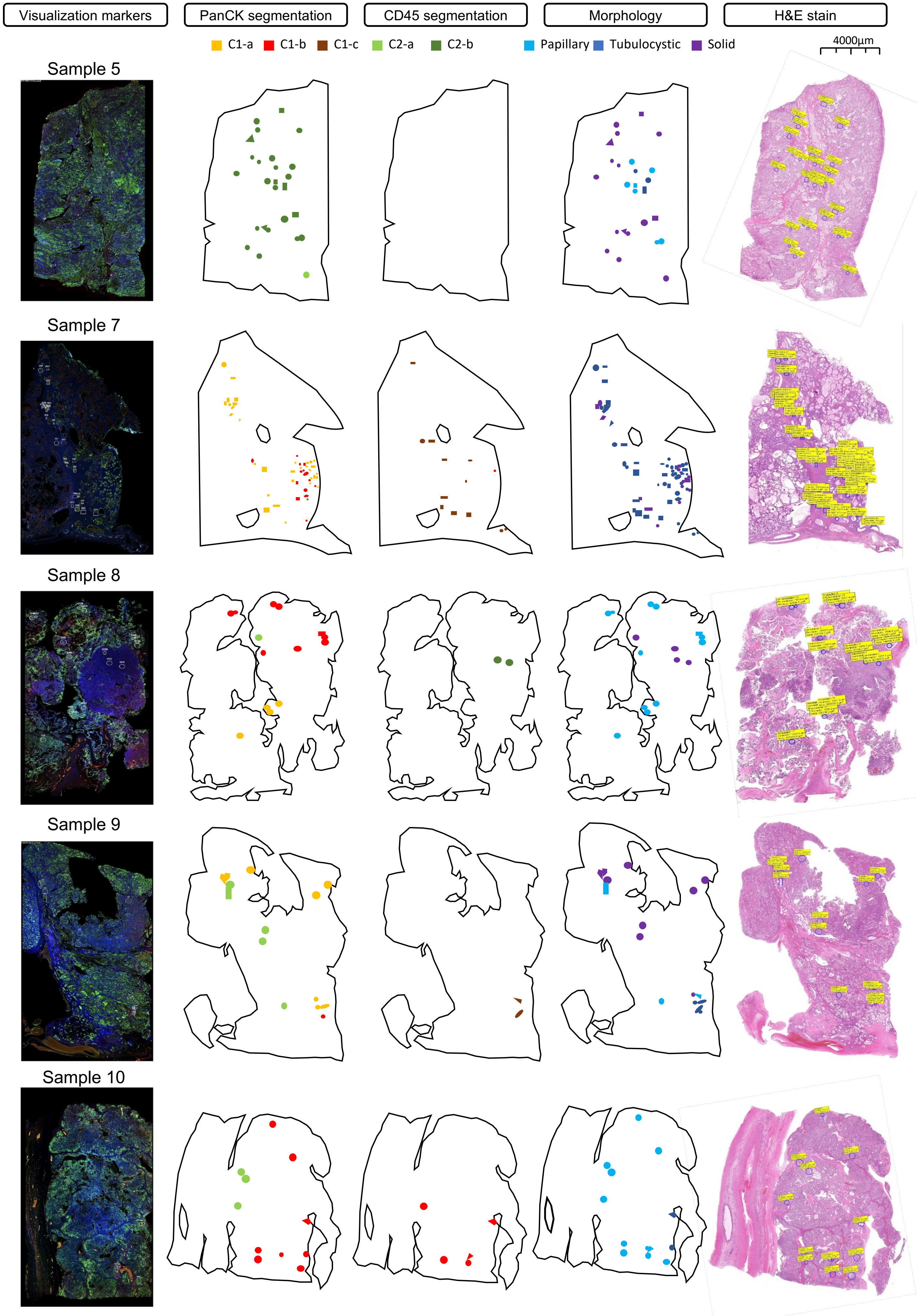

### Supplementary figure 3

#### Supplementary Figure 3

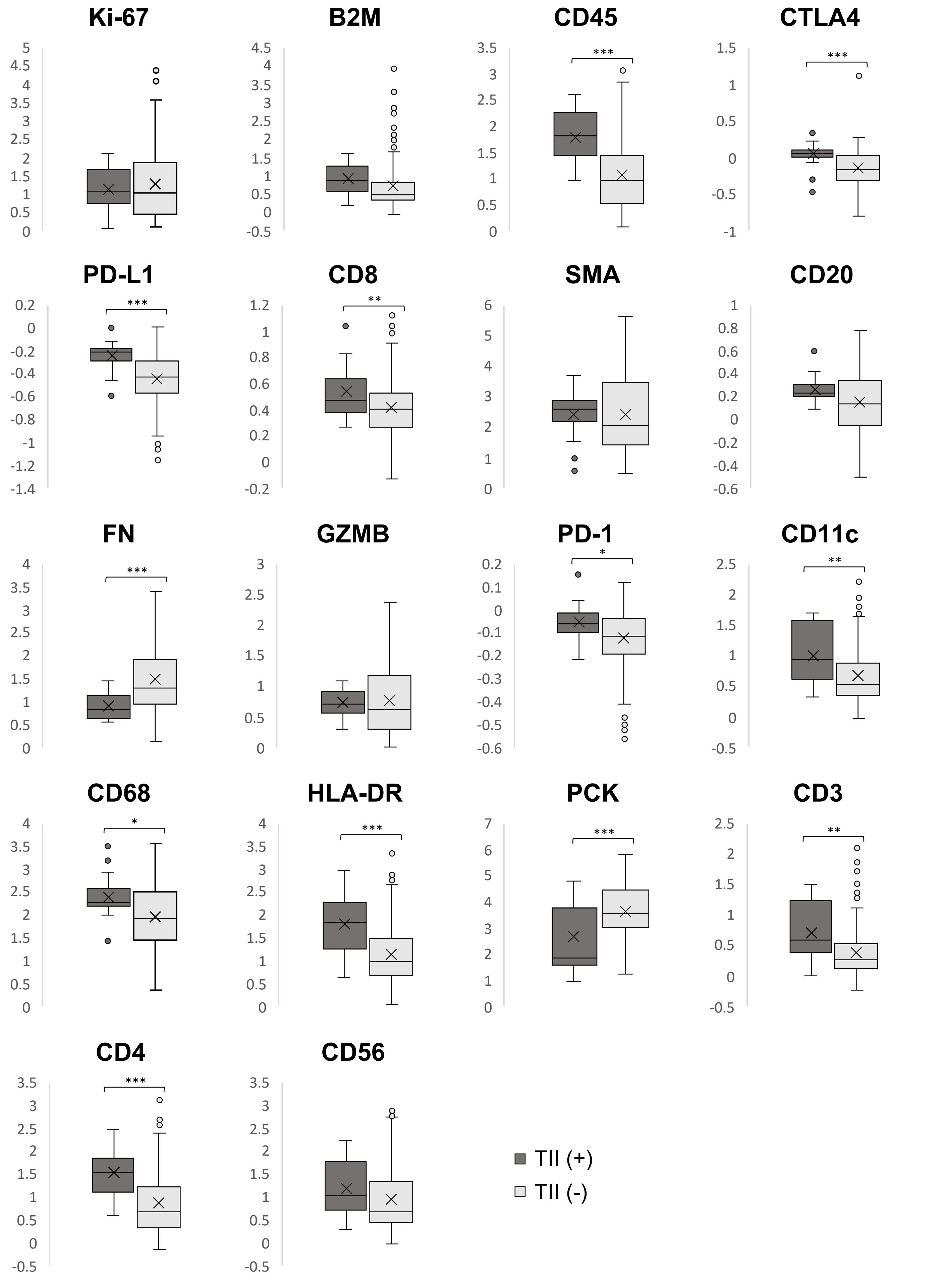

### Supplementary figure 4

#### Supplementary Figure 4

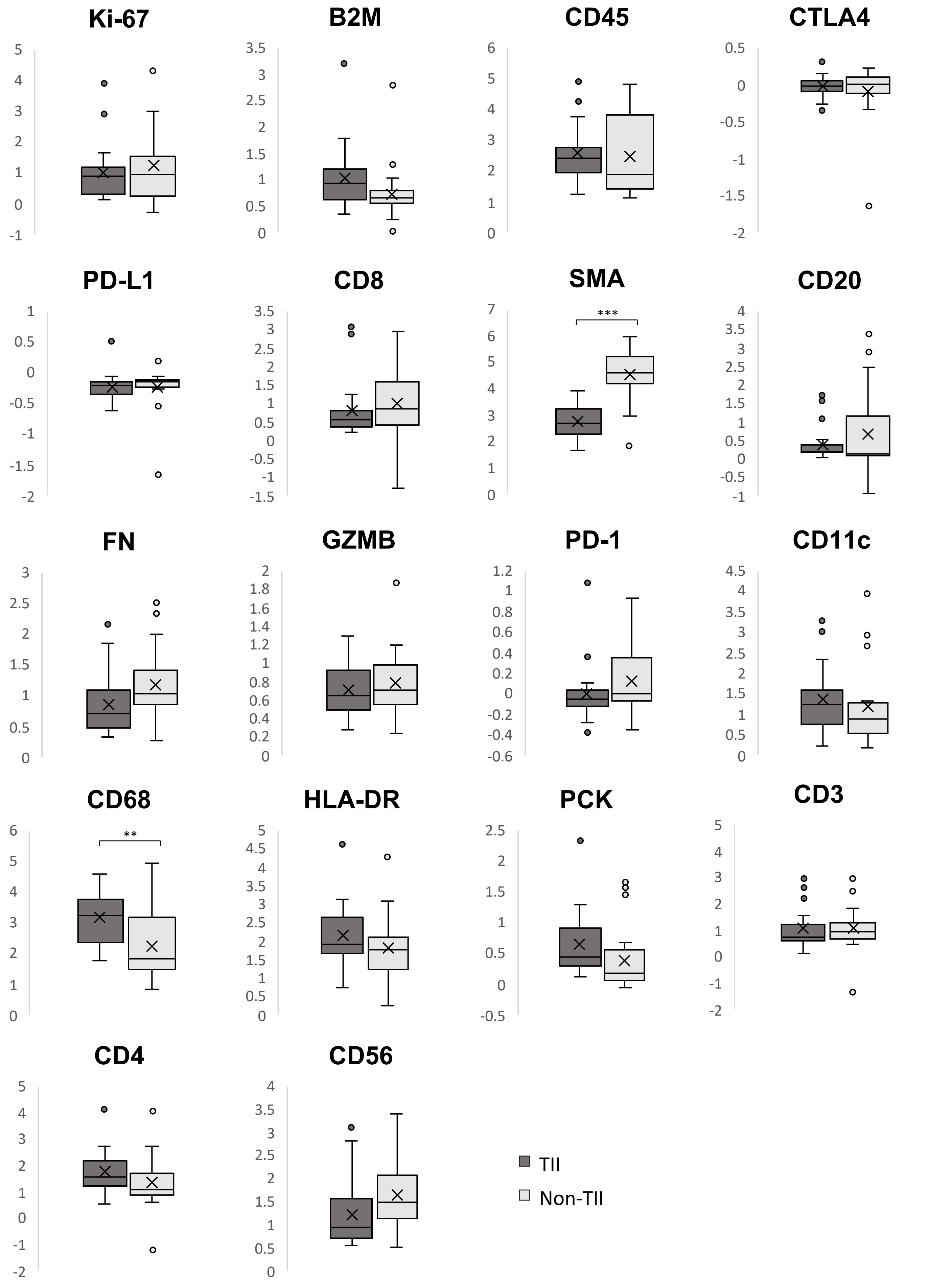

### Supplementary figure 5

##### Supplementary Figure 5

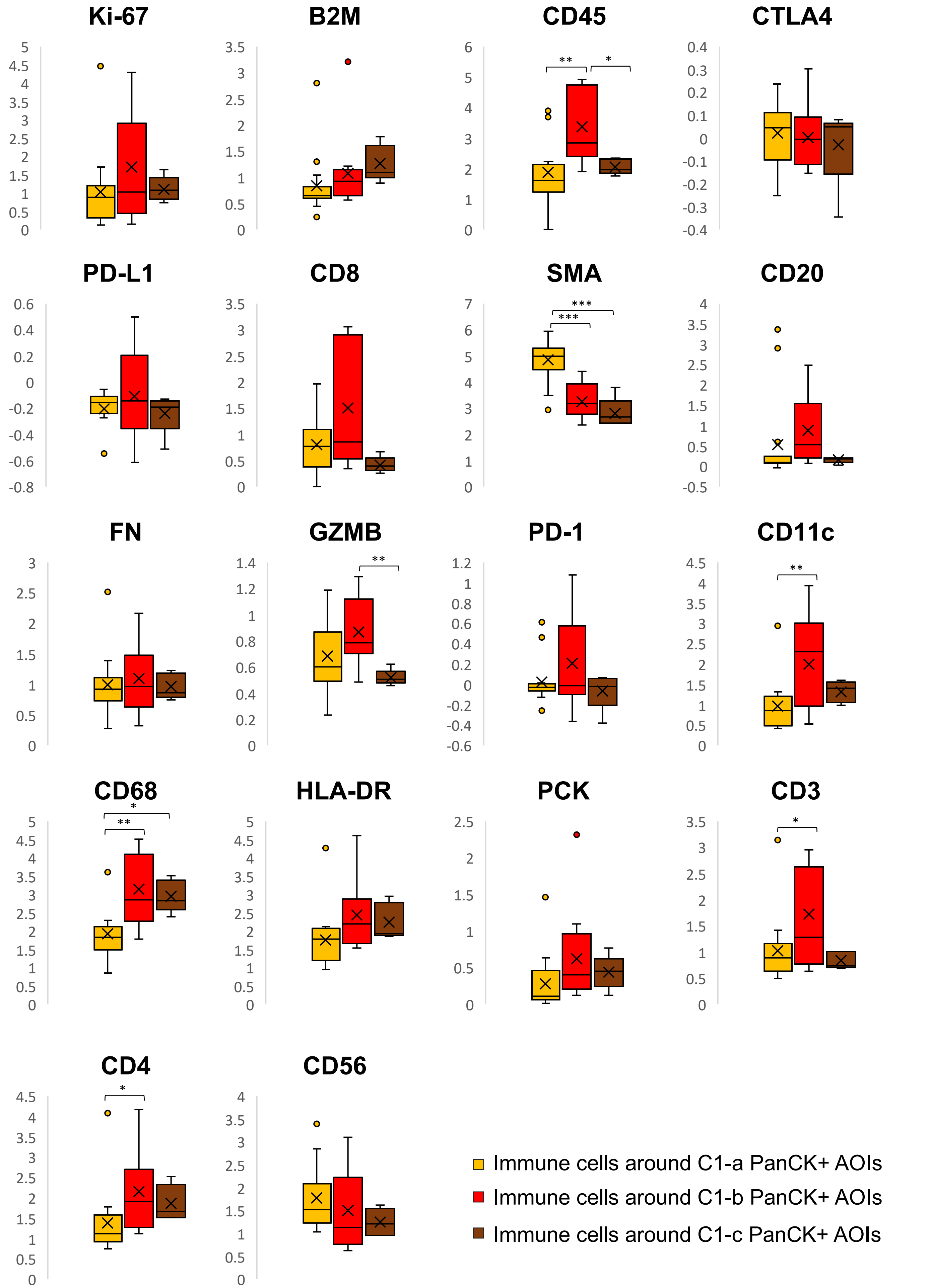

### Supplementary figure 6

#### Supplementary Figure 6

**a**

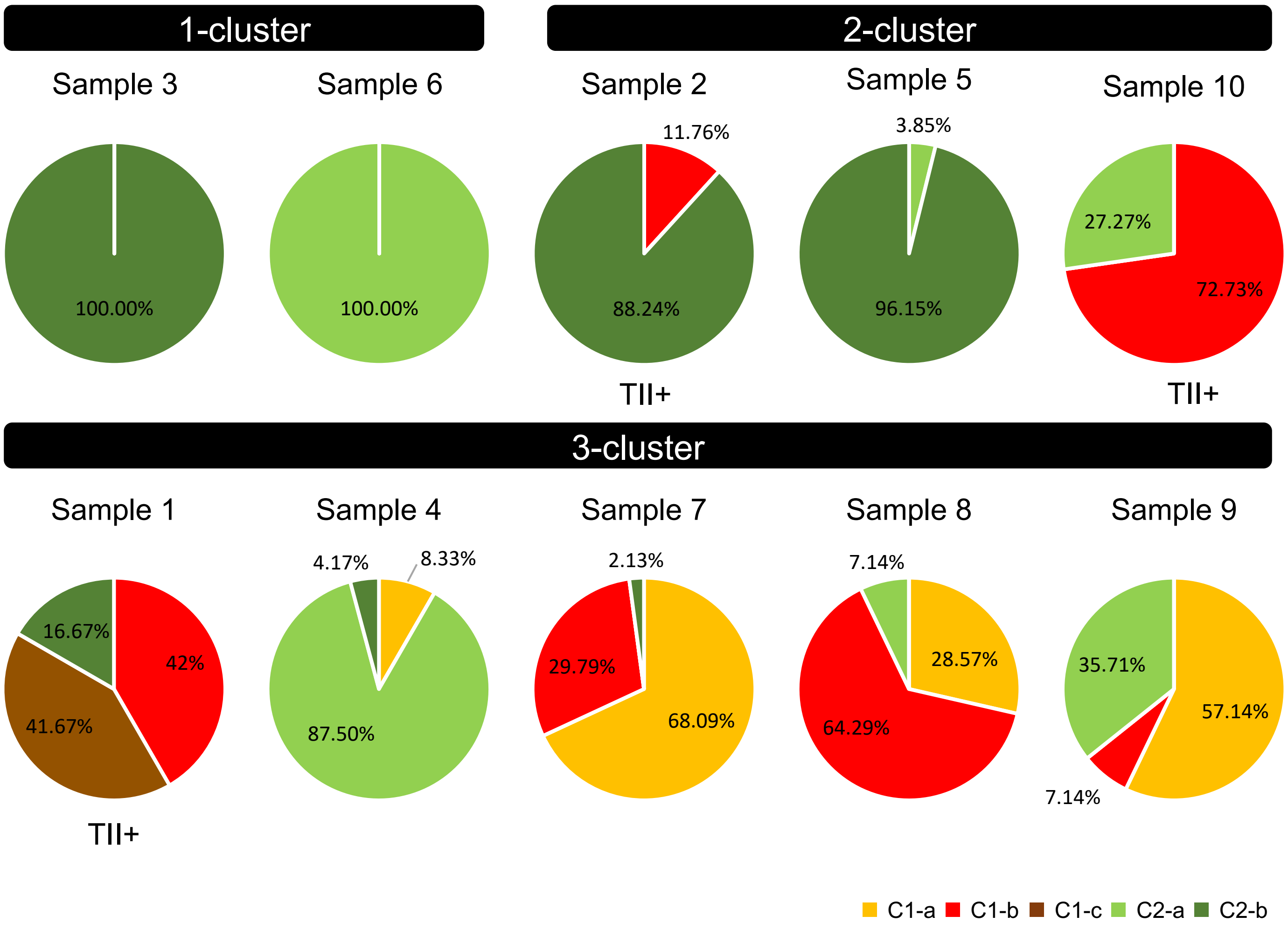


**b**

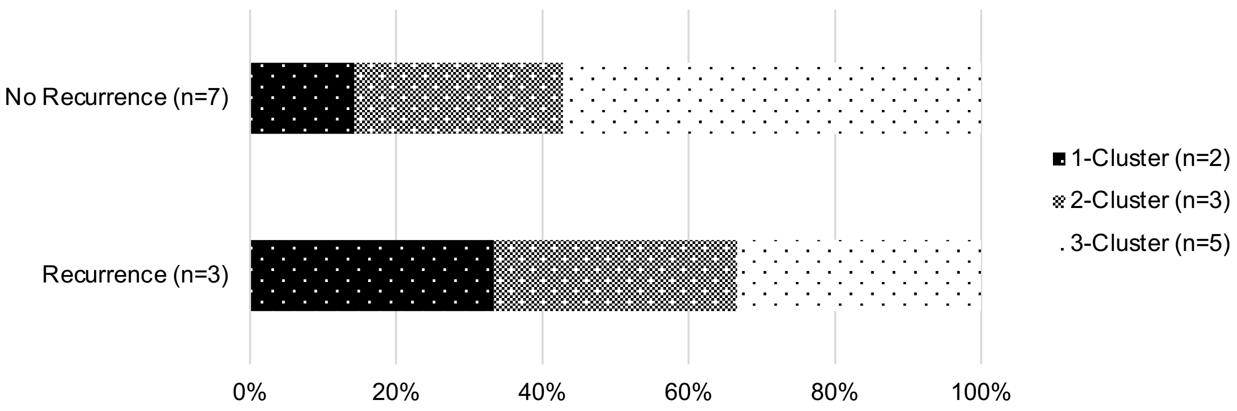

### Supplementary figure 7

#### Supplementary Figure 7

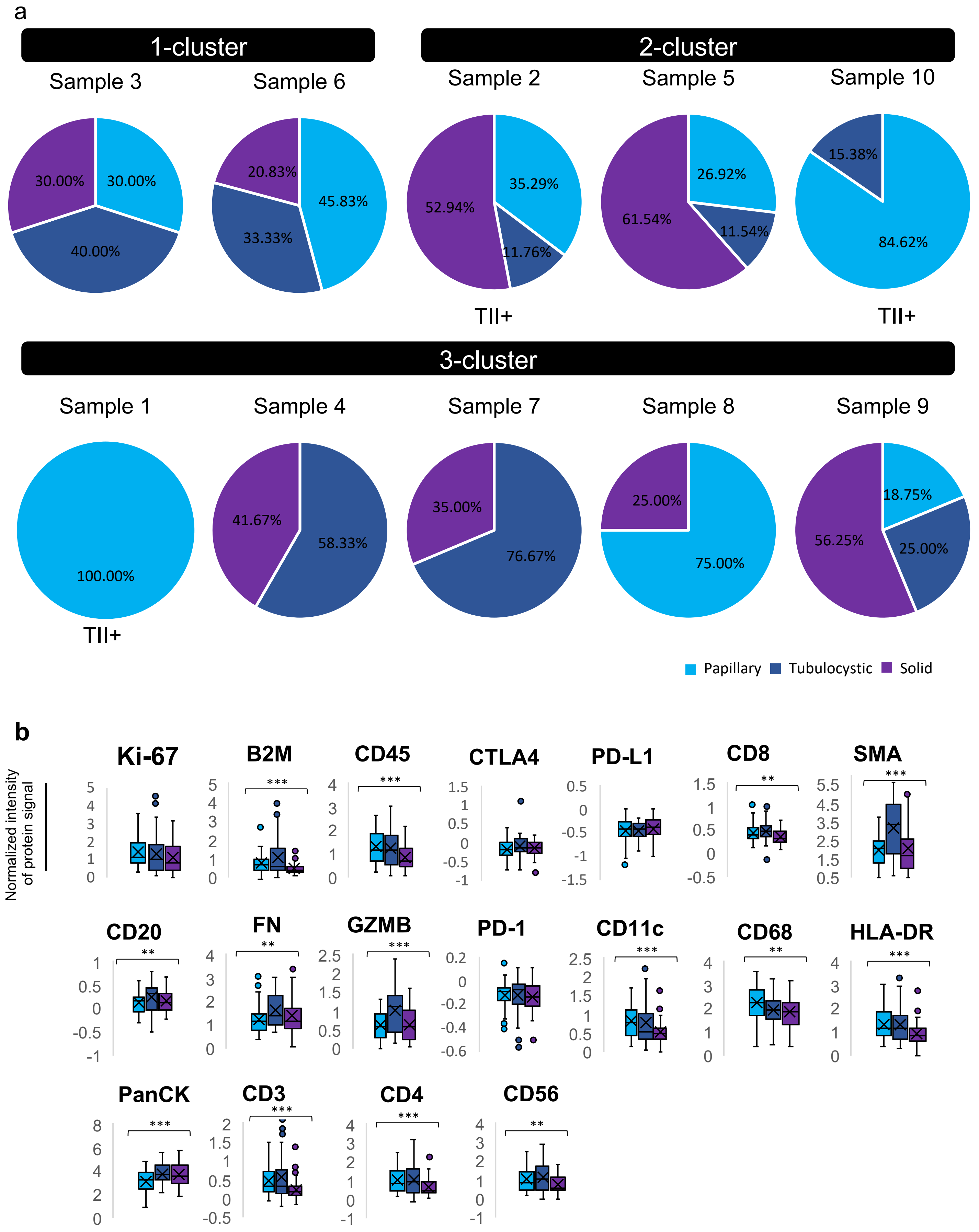
